# High-dose DFMO alters protein translation in neuroblastoma

**DOI:** 10.1101/2025.01.23.634327

**Authors:** Andrea T. Franson, Kangning Liu, Rohan Vemu, Elizabeth Scadden, Yimei Li, Annette Vu, Michael D. Hogarty

## Abstract

DFMO has been studied as a cancer therapeutic at doses ranging from 500-9,000 mg/m2/day. Lower doses are favored for cancer prevention studies while higher doses, often with chemotherapy, are studied in refractory cancers. DFMO inhibits the rate-limiting enzyme in polyamine synthesis, ornithine decarboxylase (ODC), an oncogene transcriptionally regulated by MYC. *MYC* genes are the principal oncogenic drivers of neuroblastoma, and *ODC1* is co-amplified in a subset with dismal outcome, so DFMO is a rational therapeutic candidate. Low-dose DFMO has now been FDA-approved for high-risk patients though the mechanisms for its anti-tumor activity, and the exposures required to elicit them, remain obscure. We sought to define biomarkers of activity across exposures achieved in the clinic with low through high-dose DFMO. Polyamines support protein translation by providing spermidine, which is essential to hypusinate (and activate) the elongation factor, eIF5A. Selective binding of polyamines with tRNA and rRNA provide eIF5A-independent mechanisms of translation support. We show that low-dose DFMO does not extend survival in mouse models *in vivo* nor alter translation biomarkers *in vitro*. High-dose DFMO consistently extends survival in neuroblastoma models, and, in a subset of neuroblastoma cell lines, inhibits eIF5A hypusination and global translation at achievable concentrations. However, the concentration required to engage these changes across many cell lines exceeded that achievable even with high-dose DFMO. No correlation was seen among *MYCN* and/or *ODC1* copy number and sensitivity to DFMO. Combining high-dose DFMO with additional agents to further deplete tumor polyamines may be necessary to fully engage polyamine-depletion effects on tumors, and more granular measures of translation, including codon-resolution ribosome profiling, may be required to define these effects.

**STATEMENT OF TRANSLATIONAL RELEVANCE:** Low-dose DFMO is approved by the FDA for the treatment of neuroblastoma. The depletion of tumor polyamines has been shown to have activity against tumors with activated *MYC* signaling, like neuroblastoma, yet the degree of polyamine depletion required, the mechanisms by which this impedes tumor progression, and the DFMO exposures required to enable these are poorly understood. Here we evaluate alterations in protein translation as putative mechanisms for DFMO activity. Translation biomarkers and colony formation can be inhibited by DFMO *in vitro* at exposures achievable *in vivo* with high-dose DFMO. Similarly, high-dose DFMO, but not low-dose DFMO, extends neuroblastoma-prone mouse survival. These findings support studying DFMO at higher doses and in therapeutic combinations that further augment polyamine depletion within tumors.

## INTRODUCTION

Neuroblastoma is a common and highly lethal childhood solid tumor, accounting for up to 15% of childhood cancer-related deaths [1]. While low grade and spontaneously regressing neuroblastomas occur, the majority of children have neuroblastomas with high-risk clinical or biological features. Despite treatment that includes dose-intensive chemotherapy, stem cell rescue, radiotherapy and immunotherapy, long-term survival for high-risk neuroblastoma remains poor and survivors often suffer chronic treatment-related medical conditions [1, 2]. The development of more effective and less toxic treatments that target essential tumor functions provides an opportunity to improve outcomes.

High-risk neuroblastomas frequently have genomic amplification of the *MYCN* proto-oncogene, a member of the *MYC* helix-loop-helix transcription factor family, and this amplification independently portends a poor prognosis [3, 4]. High-risk tumors without *MYCN* amplification often have *MYC* or *MYCN* activation through alternate mechanisms [3, 5, 6], including enhancer amplification or enhancer hijacking translocations [7], such that deregulated *MYC* is the principal oncogenic driver. The *MYC* family is among the most frequently deregulated in human cancer, with transcriptional activities that link the licensing of cell cycle entry with the metabolic processes needed to create biomass [8, 9]. Among these mission-critical *MYC* functions in proliferating tissues is the upregulation of protein translation machinery since proteins, tRNAs and rRNAs comprise the majority of cell biomass. Indeed, a plethora of transcriptome studies consistently identify protein synthesis, ribosome proteins, ribosome biogenesis, and polyamine synthesis as processes most highly linked to MYC, including in neuroblastoma [8, 10–12].

Direct pharmacologic inhibition of MYC has proven difficult but targeting its key oncogenic outputs provides an alternative therapeutic opportunity. Polyamines are essential oncometabolites found in nearly all living organisms and intracellular levels are tightly regulated [13]. They are abundant in proliferating cells and tissues, including cancers, and depleted from senescent, differentiated or otherwise quiescent tissues [14–16]. Polyamines are cationic chaperones that complex with rRNAs and tRNAs to support translation [17, 18]. They are also required for the post-translational modification of the eukaryotic initiation factor 5A (eIF5A) via an enzymatic modification that utilizes the polyamine spermidine to hypusinate a lysine residue to yield the active form, hypusinated-eIF5A [19]. *MYC* genes broadly regulate polyamines to support protein translation [15, 20]. In high-risk neuroblastomas, polyamine metabolism is induced by *MYCN*’s direct regulation of *ODC1*, which encodes ornithine decarboxylase, the rate-limiting enzyme in polyamine synthesis [21], further supported by transcriptional regulation of essentially all polyamine enzymes to support polyamine sufficiency [22].

*ODC1* is itself an oncogene [23, 24] that catalyzes the decarboxylation of ornithine to putrescine, the first-order polyamine that is subsequently converted to spermidine and spermine by their respective aminopropyl transferases. The activities of both ODC and the second rate-limiting enzyme in polyamine synthesis, adenosylmethionine decarboxylase (encoded by *AMD1*), have the shortest half-lives of any mammalian enzyme. ODC is further regulated by direct inhibitors (antizymes) that are themselves regulated by antizyme inhibitors [25] through frameshifting at the level of translation regulated by the polyamines themselves. Such exceptional evolutionarily conserved control over polyamine abundance underscores the critical roles they play in cellular homeostasis (reviewed in [26]). We previously identified *ODC1* amplification as co-occurring with *MYCN* amplification in a subset of neuroblastomas, the first example of co-amplification of an oncogene with its oncogenic target gene [27]. *ODC1* is the gene most frequently co-amplified with *MYCN* and portends an exceptionally poor prognosis [28]. Importantly, independent of *ODC1* amplification, high *ODC1* expression itself correlates with reduced survival [27, 29], and a 12-gene polyamine signature score was found to be highly prognostic, with incrementally poorer outcome with increasing signature expression [22].

ODC is therefore a target of interest in high-risk neuroblastoma. Difluoromethylornithine (DFMO; eflornithine) is a direct inhibitor of ODC by serving as a false substrate for the enzyme, covalently binding and inactivating ODC. DFMO has been used at high doses (12-18 gm/m^2^/day) in the treatment of sleeping sickness (*Trypanosomic* encephalitis) since ODC is an essential *Trypanosoma* gene. As polyamines are essential to support tissue proliferation, DFMO has been explored as a cancer agent, usually at or near the maximal tolerated dose. The recommended Phase 2 dose of DFMO when combined with chemotherapy is 9 gm/m^2^/day in adults [30, 31] and 6.75 gm/m^2^/day in children [32]. However, DFMO at lower doses (≤1.5 gm/m^2^/day) has been tested in the chemoprevention of epithelial cancers [33] and in neuroblastoma [34–36]. Data from the latter setting includes a non-randomized Phase 2 trial compared to a historical control arm propensity matched for risk factors. This trial led to FDA approval of low-dose DFMO for the maintenance treatment of neuroblastoma following upfront chemotherapy, stem cell rescue and immunotherapy [37].

Here, we measured biomarkers of protein translation in neuroblastoma cell lines of differing genomic status with respect to *MYCN* and *ODC1* copy number, to assess vulnerabilities to DFMO-mediated effects. We tested a range of DFMO concentrations that span exposures achievable in children receiving low-dose through high-dose DFMO regimens. Further, since DFMO has been shown to delay tumor progression and extend survival in multiple complementary mouse models of neuroblastoma as a single agent and in combination with chemotherapy [27, 38], we sought to identify the exposures required for these anti-tumor activities *in vivo*. A better understanding of DFMO’s mechanisms of anti-tumor activity and the exposures required to elicit them is necessary to fully leverage DFMO and complementary polyamine reducing therapeutics for clinical use.

## MATERIALS AND METHODS

### Cell lines and Tissue Culture

Human neuroblastoma cell lines were obtained courtesy of Dr. Garrett M. Brodeur (Children’s Hospital of Philadelphia, PA) and the Childhood Cancer Repository (cccells.org). Cells were cultured in RPMI 1640 (Life Technologies; Carlsbad, CA) supplemented with 10% fetal bovine serum, 1% L-glutamine (Corning; Corning, NY), 1% penicillin/streptomycin (Life Technologies), and 0.05% gentamicin (Life Technologies). Tissue culture was performed at 37°C in a humidified atmosphere of 5% CO_2_. All cell lines were identity-confirmed every 6 to 8 months using short tandem repeat (STR)-based genotyping (AmpFISTR, Applied Biosciences; Foster City, CA) and matched to COG cell line database (cccells.org).

### Colony Formation Assay

DFMO (α-Difluoromethylornithine, Eflornithine) was provided by Susan Gilmore (Lankenau Institute for Medical Research; Wynnewood, PA) and Patrick Woster (courtesy of ILEX Pharmaceuticals). Neuroblastoma cells were exposed to DFMO at concentrations of from 0.15-5 mM in tissue culture media for 120 hours. After exposure, 4 x 10^3^ cells were plated in triplicate in 6-well plates in RPMI without DFMO. Plates were kept at 37 °C for 14 days, with culture media changed at day 7. Colonies were fixed with acetic acid in methanol (14% v/v) for 5 minutes and stained with 0.005% crystal violet in PBS and counted using ImageJ 1.50i (imagej.com.nih.gov/ij).

### Quantitative PCR

Quantitative PCR studies were carried out on DNA isolated from neuroblastoma cell lines using the QIAmp DNA Mini Kit (Quiagen; Valencia, CA) per supplied protocol. For each well, 10 ng DNA (in 2 μL of ddH_2_O) was added to 2.5 μL nuclease-free water, 0.5 μL of primer, and 5 μL of 2x TaqMan^®^ Genotyping Master Mix (Applied Biosystems; Waltham, MA). Each sample was run in triplicate, and serial dilutions of human retinal pigment epithelial cell line (hTERT RPE1) DNA was used to create a standard curve from 0.1-20 ng of genomic DNA, respectively. Copy numbers of *MYCN* and *ODC1* were compared to *MTTP* (disomic in >90% of neuroblastoma tumors [39]). The samples were run on the 7900HT Fast Real-Time System (Applied Biosystems) with cycling as follows: 2 minutes at 50°C, 10 minutes at 95°C, followed by cycles of 15 seconds at 92°C and 60 seconds at 60°C for 40 cycles. A manual C_T_ threshold was set at 0.2.

Primers: *MTTP* assay #Hs04877005_cn, *ODC1* assay #Hs02651018_cn, and *MYCN* assay #Hs00201049_cn (Applied Biosystems), chosen for similar amplicon size (80-100 bps) for matched amplification efficiency. Analysis was performed using a simple linear regression model using the hRPE1 curve to calculate relative “dose” of each gene, expressed as a ratio to the control gene dose (*MTTP*). The *MYCN/MTTP* or *ODC1/MTTP* ratio of each sample was divided by the ratio for the control sample (hTERT-RPE1). A pre-determined threshold of 10^2^ was used to call amplification of *MYCN* or *ODC1* compared to *MTTP* and normalized to hRPE1. *MYCN* amplification is routinely defined for neuroblastoma cell lines via fluorescence in situ hybridization (FISH) analysis. *MYCN* amplification status is known on all cell lines in this study, serving as an internal comparator. Previously obtained SNP data were also analyzed and served as a confirmatory method for *ODC1* amplification status when available, using the Log2 median normalization to assess for amplification of the genes of interest (*MYCN* and *ODC1*).

### Puromycin Incorporation

Cells were treated with DFMO (0.15-5 mM in DMSO) and/or AMXT-1501 (2.5 µM; MedChem Express, Monmouth Junction, NJ), in routine tissue culture media for 3-5 days. Cells in growth phase (<80% confluence) were labeled with 1 μM puromycin at 37°C for 1 hour, then media was replaced with Versene and 5 μM cycloheximide to inhibit protein translation. Lysates were separated electrophoretically (per below) and transferred to PVDF membranes (Invitrogen), blocked with milk buffer, and detected with a mouse monoclonal puromycin antibody (clone 12D10, #MABE343; Millipore) at 1:10,000 at 4°C overnight, followed by goat anti-mouse secondary at 1:3000 dilution. Densitometry was performed using ImageJ.

### Isoelectric Focus

The isoelectric focusing assay allows for the separation of proteins according to their isoelectric point (pI) within a fixed pH gradient within a polyacrylamide gel [40]. Protein lysates were prepared from cells treated with DFMO for 5 days (0.15-5 mM) in 2X sample buffer (Novex IEF, LC5371, Invitrogen; Waltham, MA) without boiling or loading dye, loaded into an isoelectric focusing gel pH 3-7 (Novex, EC66452, Invitrogen) and run per manufacturer’s instructions with a pH marker lane (Novex IEF Marker 3-10, Serva Liquid Mix, 39212.01, Invitrogen) and transferred to a PVDF membrane using the iBlot (Invitrogen) system for 8 minutes at setting P3, placed in 5% skim milk blocking buffer in TTBS (w/v) x1 hour, and blotted for eIF5A at 1:10,000 dilution.

### Immunoblots

Cells were treated with DFMO (0.15-5 mM), AMXT-1501 (2.5 µM), and/or MLN0128 (100 µM; MedChem Express). Protein lysates (20 µg) were run on a 4-12% Bis-Tris NuPAGE Gel at 150V for 90 min on ice. Proteins were transferred to PVDF membranes and blocked for 60 min with a 5% milk/TTBS solution, followed by a TTBS wash. Membranes were incubated overnight at 4°C with anti-hypusine (1:3000; rabbit polyclonal, Millipore ABS1064), anti-puromycin (1:5000; mouse monoclonal, Millipore MABE343), anti-ODC1 (1:5000; mouse monoclonal, Novus 32887), anti-β-tubulin (1:3000; mouse monoclonal, Millipore T8328), anti-eIF5A (1:3000; mouse monoclonal, BD Transduction 611976), anti-GAPDH (1:1000; rabbit polyclonal, Santa Cruz SC-25778), anti-phospho-4EBP1 (1:1000, rabbit polyclonal, Cell Signaling Technology 9451), and anti-4EBP1 (1:1000, rabbit monoclonal, Cell Signaling Technology 9644) antibodies. Membranes were washed three times for 5 min with TTBS and incubated with anti-mouse (1:3000, Invitrogen) or anti-rabbit secondary antibody (1:3000, Invitrogen) for 60 min at RT, washed again three times with TTBS and developed with HRP chemiluminescent substrate (Immobilon ECL Ultra Western HRP Substrate, Millipore). Densitometry with ImageJ was used to assess.

### *TH-MYCN* studies

129×1/SvJ mice transgenic for the *TH-MYCN* construct [41] were originally provided by Bill Weiss (University of California, San Francisco, San Francisco, CA). All murine studies were approved by the Institutional Animal Care and Utilization Committee at The Children’s Hospital of Philadelphia (Philadelphia, PA). *TH-MYCN*^+/-^ mice were bred and offspring genotyped from tail-snip–isolated DNA using qPCR [27]. *TH-MYCN*^+/+^ mice have fully penetrant lethal neuroblastomas. *TH-MYCN*^+/+^ pups were randomized to *ad libitum* water (control), or water supplemented with 0.25%, 0.5% or 1% DFMO from birth onward (given to mothers pre-wean and to pups directly post-wean). Mice were monitored by a single animal technician and sacrificed for signs of symptomatic tumor progression to define survival time. To estimate DFMO exposure, water intake for a group of 4-6 mice from each treatment arm was measured daily over 14 days and averaged per mouse.

### Statistical analyses

GraphPad Prism version 10.2.2 (San Diego, CA) was used for all statistical analysis. All quantitative data were presented as the mean +/- standard deviation (SD) as indicated of at least three independent experiments. One-way analysis of variance (ANOVA) followed by Dennett’s post hoc multiple comparison test was used to determine the statistical significance of treatment groups versus control. Welch’s t-test was used to compare cell doubling time in *MYCN*-amplified versus *MYCN* non-amplified cell lines. Statistical significance was defined as p < 0.05 (*), p < 0.01 (**), p < 0.001 (***).

## RESULTS

### Co-amplification of *ODC1* with *MYCN* in neuroblastoma

*MYCN* amplification occurs in ∼20% of neuroblastomas overall and ∼40% of high-risk tumors [42]. Co-amplification of *ALK* (2p23.2; 13.5 Mb centromeric to *MYCN*) occurs in 3-5% of tumors while isolated *ALK* amplification is rare [43–45]. Co-amplification of *ODC1* (2p25.1; 5.5 Mb telomeric to *MYCN*) with *MYCN* has not been well characterized so we reviewed publicly available databases and the literature and assessed copy number at these genes across a panel of neuroblastoma cell lines. We identified 12 publications and datasets that defined *MYCN* and *ODC1* status for >1,200 primary neuroblastomas and >60 cell lines utilizing a range of methodologies (array-CGH, SNP-array, NGS; **Table 1**). We excluded earlier reports using Southern blot methodology [46–48]. No tumor was identified with *ODC1* amplification in the absence of *MYCN* amplification. The proportion of primary neuroblastomas with *ODC1* amplification was 3.7%, though these are not unselected cohorts as evidenced by their range of *MYCN* amplification from 14-49%. When restricted to *MYCN* amplified tumors, *ODC1* is co-amplified in 12.6%. A higher proportion of neuroblastoma-derived cell lines carry mutations in oncogenes that provide a tissue culture growth advantage, as with *MYCN* amplification. In 69 neuroblastoma cell lines, *ODC1* was co-amplified with *MYCN* in 14%, relatively neutral with respect to tissue culture growth with concurrent *MYCN* amplification. Amplicons often carry passenger genes in immediate proximity to the gene providing the selective advantage. In neuroblastoma, *DDX1*, *NBAS* and *FAM49A* are frequently co-amplified with *MYCN* on a contiguous 1-3 Mb amplicon [49]. In contrast, *ODC1* (and *ALK*) are more distant and are discretely amplified, as evidenced by their genomic pattern on SNP-arrays (**Fig.1**).

**Figure 1.**
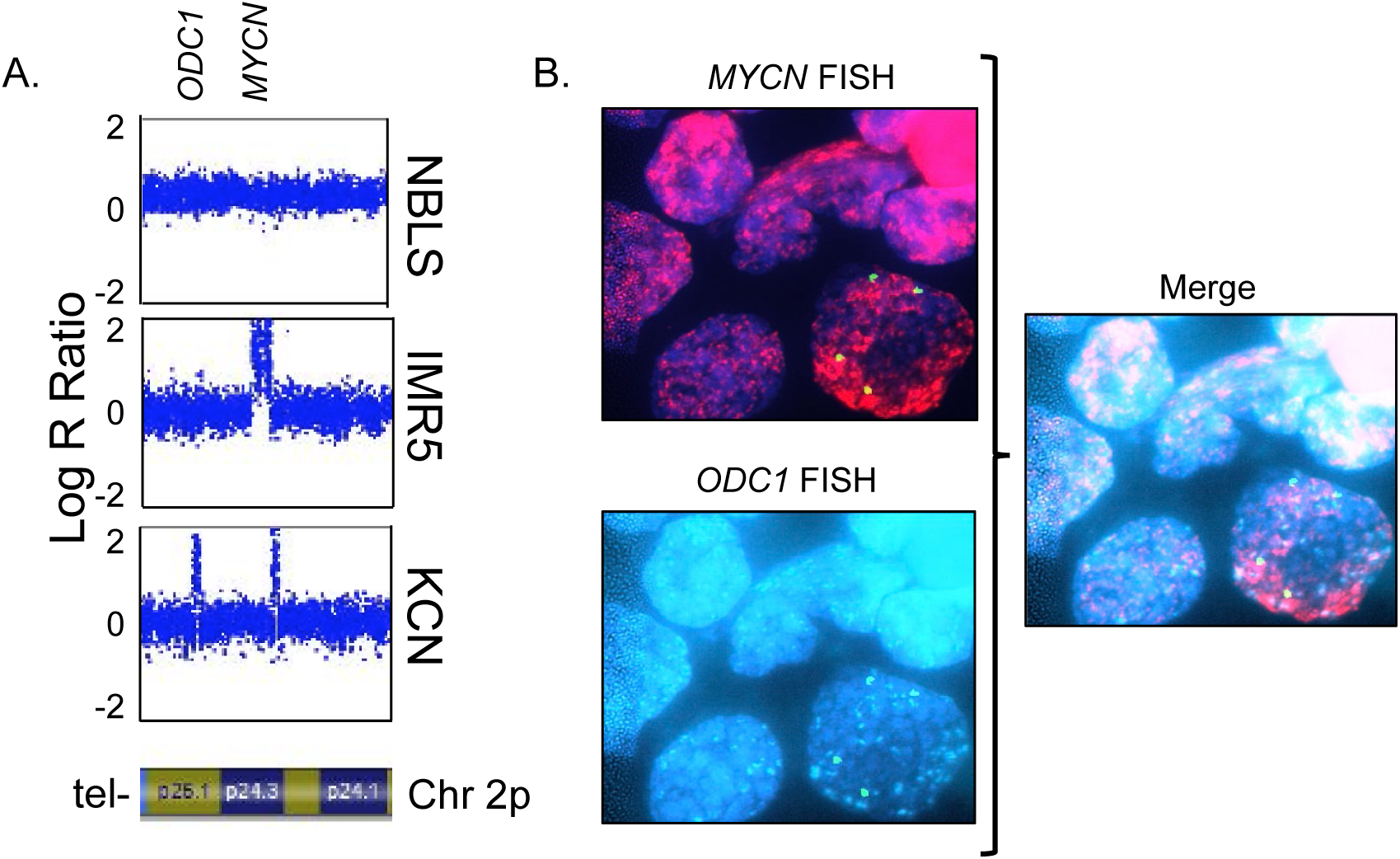
*ODC1* and *MYCN* co-occurring amplification. (A) Neuroblastoma cell lines with distinct genomic profiles are shown: NBLS cells have neither *MYCN* nor *ODC1* amplification, IMR5 has *MYCN* amplification (and *ALK* amplification, not shown), and KCN has both *MYCN* and *ODC1* amplification; data represents Log R ratio output from Illumina Bead-Chip SNP arrays. (B) Fluorescence in situ hybridization (FISH) results for a primary neuroblastoma sample using probes for *MYCN* (2p24.3, red), *ODC1* (2p25.1, aqua) and 2p centromere (CEN2, green) showing 4 CEN2 signals and amplification of both *MYCN* and *ODC1*. Images courtesy of CHOP Division of Genomic Diagnostics (DGD).

**Table 1.**
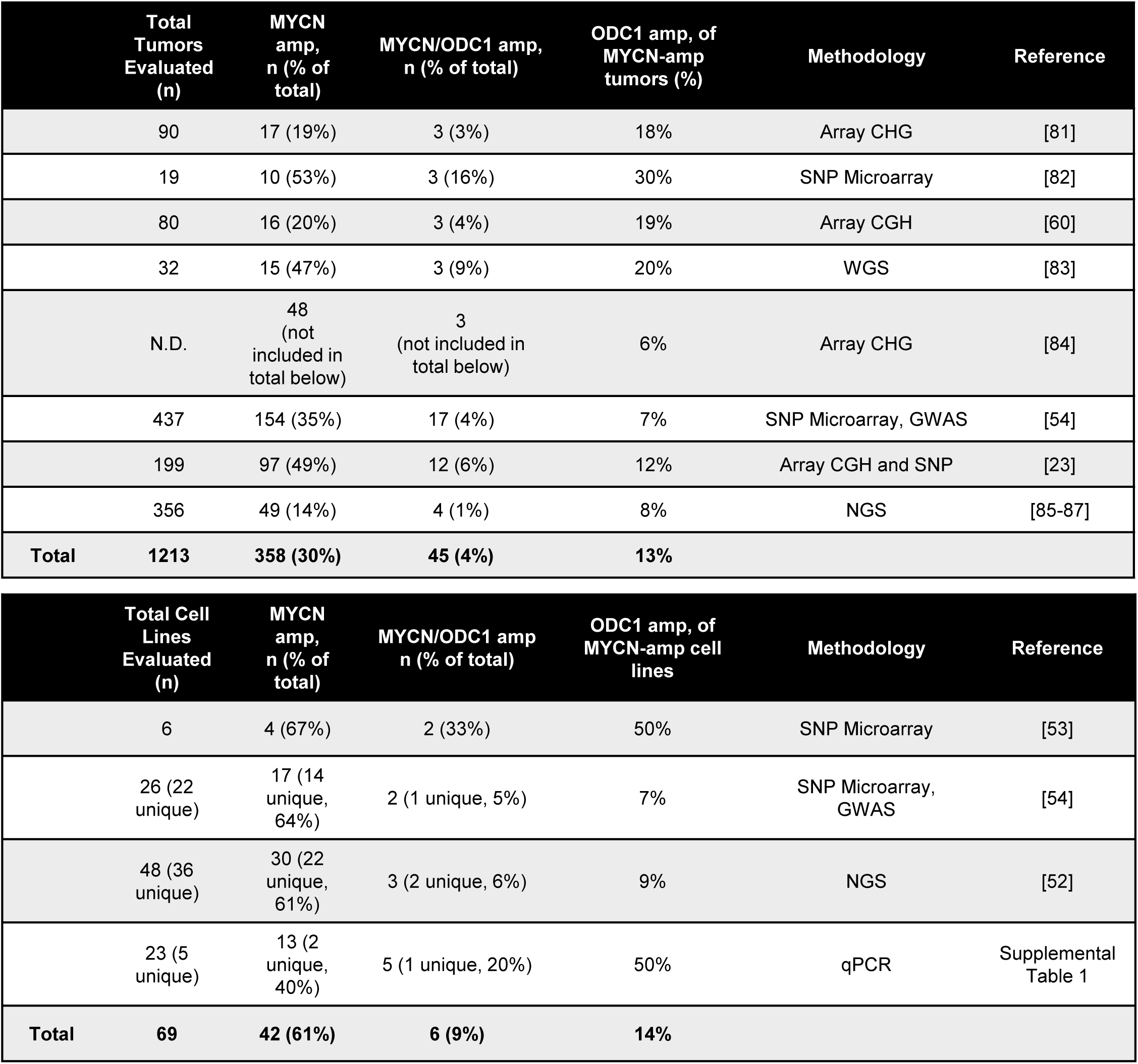
*MYCN* and *ODC* amplification in primary neuroblastomas.

We performed quantitative-PCR (qPCR) to determine the relative copy number of *ODC1* and *MYCN* in a panel of 22 neuroblastoma cell lines (21 from unique patients; one with a patient-matched post-relapse cell line [CHLA122/CHLA136]; **Supplementary Tables 1, 2**). We selected the *MTTP* gene (4q23) as a control for *MYCN* and *ODC1* copy number comparison as 4q23 is disomic in >90% of tumors and cell lines in which SNP-array data were available [50]. *MYCN* amplification was detected by qPCR in 12 of 21 unique cell lines (57%) and was concordant in both diagnosis/relapse pairs, as expected [51]. Further, *MYCN* status by qPCR was concordant with available FISH results (13 of 13 scored as amplified; 10 of 10 non-amplified). *ODC1* amplification was detected in 3 of 21 neuroblastoma cell lines overall (14%), all with *MYCN* co-amplification (3 of 12 *MYCN* amplified cell lines; 25%). SNP-array data suitable for copy number determination at these loci were available for 17 and *ODC1* copy number was concordant between SNP-array and qPCR assay (3 of 3 scored as amplified; 14 of 14 non-amplified). Concordant results were also found in comparison with orthogonal genomic datasets reported independently for many of these cell lines [52–54] (**Supplementary Table 1**).

### Clinically achievable DFMO concentrations from low to high dose in children, adults and mice and correlation with preclinical activity

To assess neuroblastoma cell line responses to DFMO *in vitro*, we first defined clinically relevant exposures. Orally administered DFMO has been studied in clinical trials for cancer at doses ranging from 500 mg/m^2^/day through >9 gm/m^2^/day, alone and in combination with chemotherapy. Pharmacokinetic studies show oral DFMO is 55% bioavailable but with significant interpatient heterogeneity [55]. C_max_, C_min,_ and AUC increase linearly through ∼4 gm/m^2^/day and above this the increase is non-linear. The FDA-approved DFMO dose for neuroblastoma maintenance therapy ranges from 0.75-1.5 gm/m^2^/day divided BID (varies by patient BSA due to tablet size), providing a measured mean C_max_ of 50 ± 22 µM [35] and estimated C_min_ of 12-20 µM at the 1.5 gm/m^2^/day dose, consistent with PK data in adults [56]. A Phase 1 trial for children with relapsed or refractory high-risk neuroblastoma studied dose-escalated DFMO with chemotherapy [32]. DFMO trough levels (C_min_) at the recommended Phase 2 dose of 6.75 mg/m^2^/day divided TID were 74 ± 60 µM with an estimated C_max_ of 150-225 µM [32], also consistent with levels obtained in adult cancer trials [57].

Much of the preclinical neuroblastoma data supporting anti-tumor activity for DFMO were obtained in genetically-engineered mice (*TH-MYCN* model; [58]) and xenografts of human-derived cell lines or PDXs. *Ad libitum* intake of DFMO in drinking water has been favored over repeated oral gavage for chronic therapy. A range from 0.5%-2% DFMO has been studied [22, 27, 59, 60]. Fozard et al measured DFMO intake at ∼1,000 mg/kg-bwt/day per mouse drinking 0.5% DFMO in water, and ∼3,000 mg/kg-bwt/day per mouse taking 2% DFMO in water [61, 62]. Consistent with this, we measured daily intake of *TH-MYCN* mice receiving DFMO in water *ad libitum* over 14 days (n=4-6 mice each at 0.25%, 0.5%, 1% and 1.5% DFMO), with intake measured at ∼550 mg/kg-bwt/day DFMO at 0.25%; 1,100 mg/kg-bwt/day DFMO at 0.5%; 2,200 mg/kg-bwt/day DFMO at 1% and 3,300 mg/kg-bwt/day DFMO at 1.5%. FDA guidelines recommend extrapolating from animal to human dose through BSA normalization. Using *K_m_*=3 for mice and *K_m_*=25 for children [63], this allometrically scales to human equivalent doses (HED) of ∼1.65 gm/m^2^/day, 3.3 gm/m^2^/day, 6.6 gm/m^2^/day and 9.9 gm/m^2^/day for 0.25%, 0.5%, 1% and 1.5% DFMO, respectively, spanning the clinically studied dose range in children.

*TH-MYCN*^+/+^ mice develop fully penetrant neuroblastoma with early lethality from tumor progression. Treating such mice with 1% DFMO does not reduce tumor penetrance as all mice die with tumor progression, but latency and survival are significantly extended [22, 60]. To assess the impact of DFMO across a range of exposures, we randomized *TH-MYCN*^+/+^ mice at birth to receive *ad libitum* water that included 0% (control), 0.25%, 0.5% or 1% DFMO (n=20-25 mice each). All mice eventually had lethal tumor progression, and mice given DFMO at 0.25% or 0.5% *ad libitum* had no change in tumor latency or survival compared to control mice (p=0.80 and p=0.31, respectively; **Fig.2**). Mice taking 1% DFMO *ad libitum* had extended survival (p<0.0001 compared to control; p<0.001 to other arms). DFMO trough levels were estimated in this model previously by taking blood for PK analyses at the end of their diurnal period of inactivity [32]. The mean DFMO “trough” (C_min_) for mice taking 1% DFMO *ad libitum* was 53.8 ± 49.5 µM, in line with human PK troughs near the HED for 1% DFMO noted above (74 ± 60 µM at 6.75 gm/m^2^/day), and similarly reflecting significant interpatient variability. We did not estimate a C_max_ given the variable intake of water and drug over time using *ad libitum* oral delivery.

**Figure 2.**
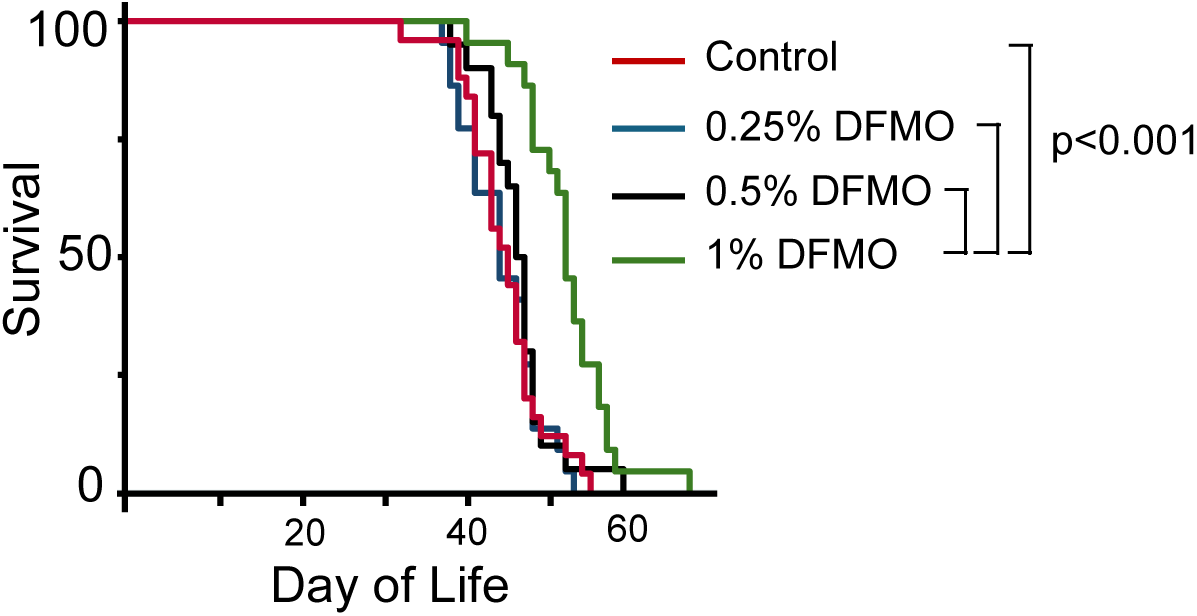
Higher DFMO exposures are required to extend life in the *TH-MYCN* neuroblastoma model. *TH-MYCN*^+/+^ mice were randomized to ad libitum water (control) or water with 0.25%, 0.5% or 1% DFMO added, fron the time of birth onward. Mice taking 1% DFMO had an extended overall survival compared to those taking 0.5% DFMO or less; p<0.001 by log-rank test; n=20-25 mice/arm. Control versus 0.25% DFMO, p=0.80; control versus 0.5% DFMO, p=0.31).

### Concentration-dependent effects of DFMO on translation biomarkers

Polyamines bind with specificity to tRNAs and rRNAs [18], their depletion from *in vitro* systems attenuates translation [64], and their depletion from cells arrests translation and proliferation [65, 66]. Yet despite their essentiality for translation, mechanistic details remain poorly characterized. Cationic chaperone-like activities of polyamines are proposed to support mTOR-mediated phosphorylation of 4EBP1, releasing eIF4E to initiate cap-dependent translation [67]. Additionally, the polyamine spermidine is required for the covalent modification that activates translation factor eIF5A via two-step hypusination (eIF5A is the only mammalian protein so modified, and spermidine is essential; [68, 69]). We therefore assessed 4EBP1 phosphorylation and eIF5A hypusination as biomarkers of DFMO activity across genomically distinct neuroblastoma cell lines. To assess global protein translation, we measured puromycin incorporation.

We exposed neuroblastoma cell lines to 5 mM DFMO and assessed 4EBP1 phosphorylation. All cell lines demonstrated multiple phospho-isoforms of 4EBP1 that were unchanged by DFMO. As a control, the mTOR1/2 inhibitor MLN0128 (sapanisertib) at 100 µM reduced some or all 4EBP1 phospho-isoforms in all cell lines tested (**Fig.3**). As DFMO at supratherapeutic concentration did not alter 4EBP1 phosphorylation, we did not test lower concentrations. We next assessed eIF5A hypusination with a hypusine-specific antibody. Only IMR5 cells had reduced hypusine content with DFMO exposures at ≥300 µM (**Fig.3B**), the remainder required up to 5 mM for this effect (data not shown). The sensitivity for detecting reduced hypusination by hypusine-specific immunoblotting is modest so we next used isoelectric separation. Incomplete eIF5A hypusination is detected as a discrete non-hypusinated band due to its distinct isoelectric point. All but one neuroblastoma cell line showed incomplete hypusination as noted by the appearance of the deoxyhypusine-eIF5A band at pI ∼5.2 (**Fig.3C**). Only CHLA136 (*MYCN* and *ODC* amplified) maintained completely hypusinated eIF5A even at supratherapeutic DFMO exposure, and CHLA15 (*MYC* amplified) had modest incomplete hypusination. In contrast, the most sensitive cell lines were IMR5 (*MYCN* amplified) and SK-N-BE2C (*MYCN* amplified). We assessed their response to DFMO across a range of concentrations. IMR5 cells were unable to fully hypusinate eIF5A at 300 µM DFMO and maximal effect was apparent by 500 µM; SK-N-BE2C cells showed incomplete hypusination at 500 µM (**Fig.3D**). MLN0128 did not impede hypusination in any cell line tested, as expected (data not shown).

**Figure 3.**
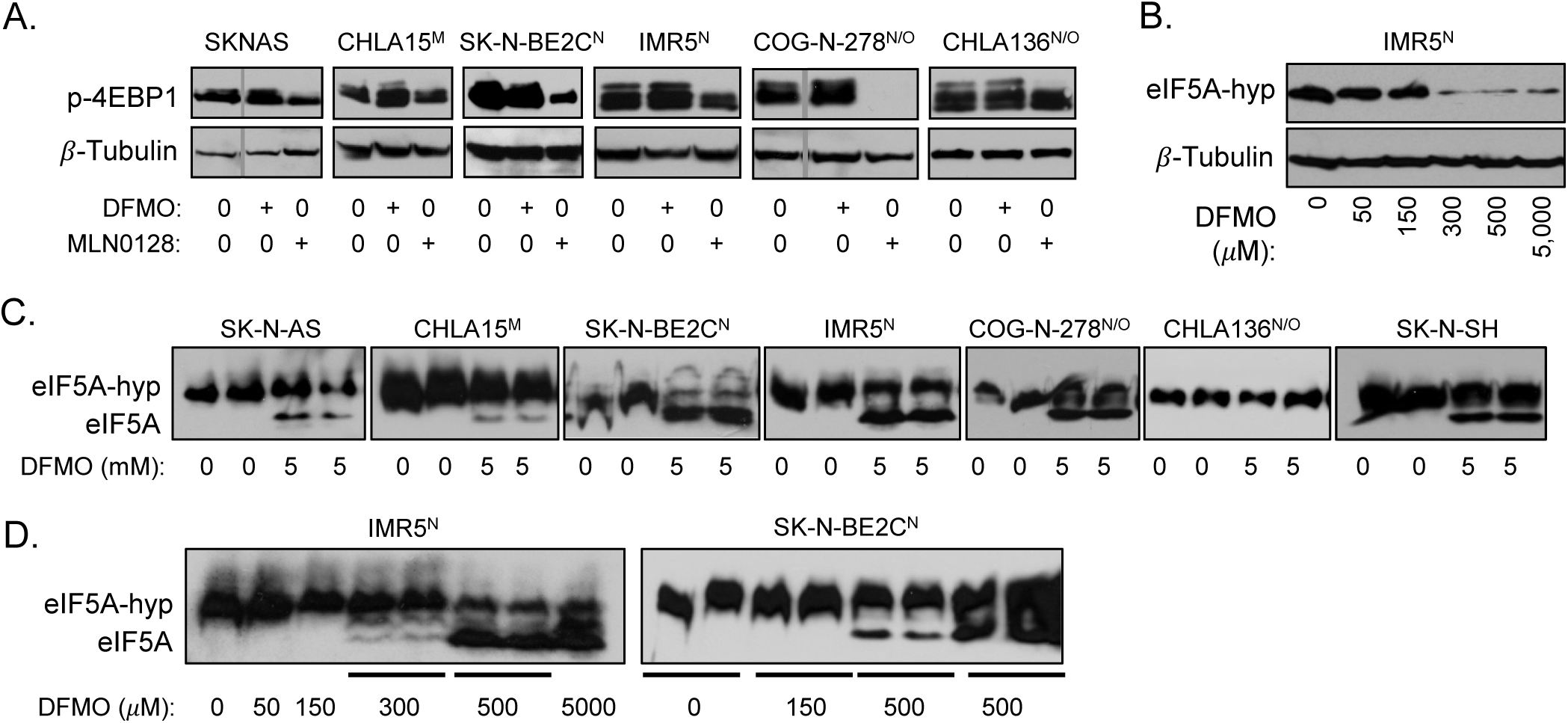
DFMO-mediated translation effects. (A) Phospho-4EBP1 is not altered by DFMO (5 uM) but is reduced by the mTOR1/2 inhibitor, MLN0128 (100 μM). (B) eIF5A hypusination is inhibited at ≥300 μM DFMO in IMR5 cells as detected using a hypusine-specific antibody; (C) more sensitive IEF immunoblot confirms shows incomplete hypusination in most cell lines at higher DFMO exposures; (D) dose response to DFMO for IMR5 and SK-N-BE2C cells (cells with higher proportion of non-hypusinated eIF5A in (C). Cell line genomic status: ^M^, *MYC* amplification; ^N^, *MYCN* amplification; ^O^, *ODC1* amplification.

### Effect of DFMO on global translation

Hypusinated eIF5A’s principal function is to resolve ribosome stalling on mRNAs encoding polyprolines and related motifs containing rigid cyclic structures [68]. However, eIF5A also broadly supports translation initiation, elongation, and termination [70]. Further, polyamines have eIF5A independent activities that promote efficient translation [65, 67]. We therefore sought to measure global translation in response to DFMO exposure. We used the puromycin incorporation assay and first aimed to determine if *MYCN* and *ODC1* amplification status was correlated with basal protein translation [71]. Puromycin incorporation, a biomarker for translation, was variable across the cell lines and no correlation was noted with amplification status or with cell line doubling-time (**Supplementary Fig.1**). Neuroblastoma cell lines had variably reduced puromycin incorporation under DFMO stress (**Fig.4**). SK-N-SH and CHLA20 had no significant reduction of puromycin incorporation at any DFMO exposure. In contrast, SK-N-AS, SK-N-BE2C, and IMR5 cells had >50% reduction in at least one replicate, and effects were most notable at concentrations >150 µM. DFMO inhibits ODC to block polyamine synthesis but compensatory polyamine uptake from the culture media can rescue this deficit (**Fig.4A, 4D, 4E**). AMXT-1501 is an inhibitor of the polyamine transporter and when combined with DFMO led to a more marked inhibition of puromycin incorporation, supporting enhanced polyamine depletion when compensatory uptake is inhibited, even in the relatively resistant SK-N-SH and CHLA20 cells (**Fig.5**).

**Figure 4.**
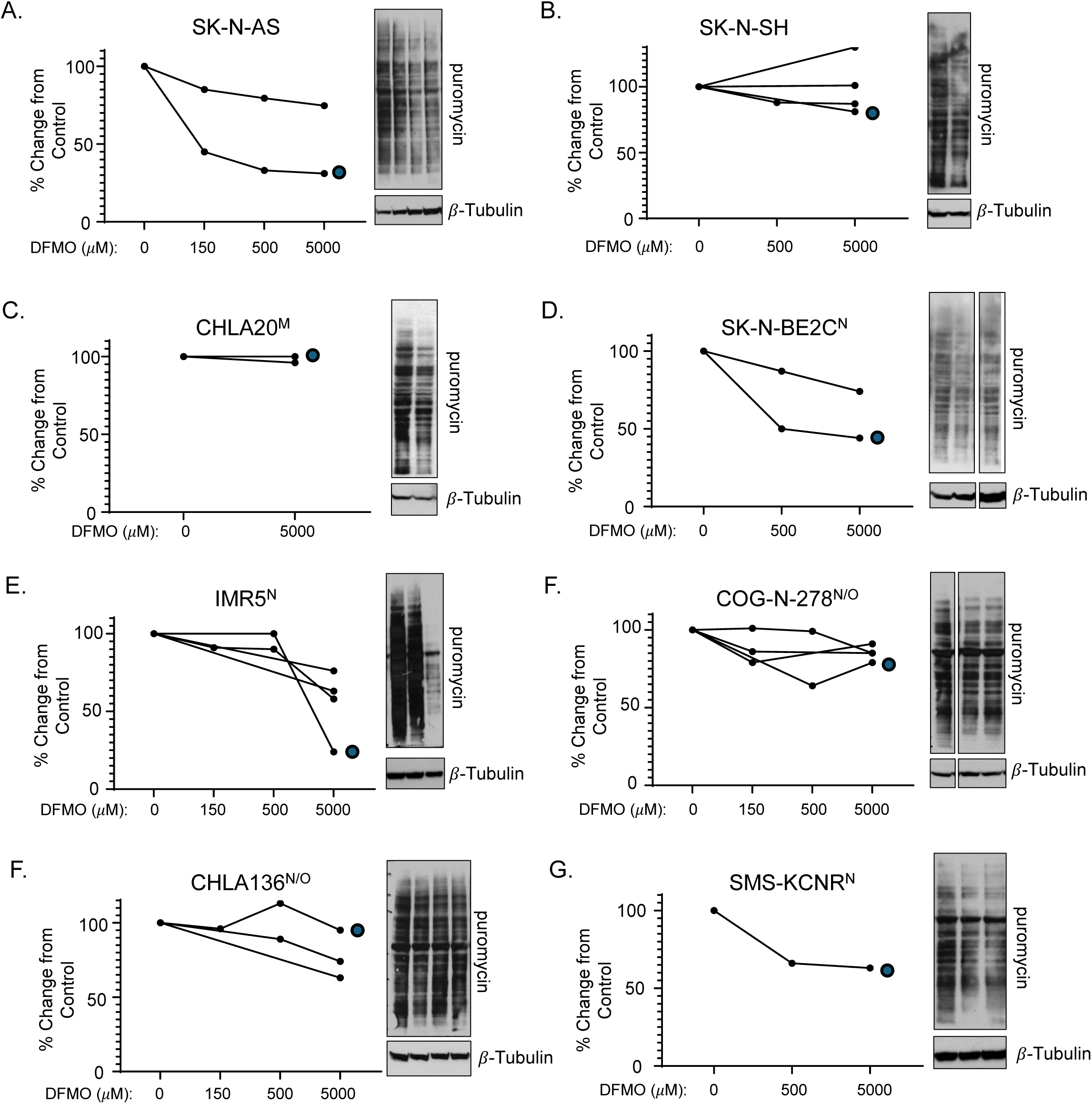
Puromycin incorporation by DFMO exposure in vitro. Puromycin incorporation was assessed across neuroblastoma cell lines of distinct *MYCN* and *ODC1* gene status, as denoted. Densitometry was used to define relative change from vehicle control lanes; replicates are shown in line graph form with the immunoblots corresponding to the data from the circle-marked line. Cell line genomic status: ^M^, *MYC* amplification; ^N^, *MYCN* amplification; ^O^, *ODC1* amplification.

**Figure 5.**
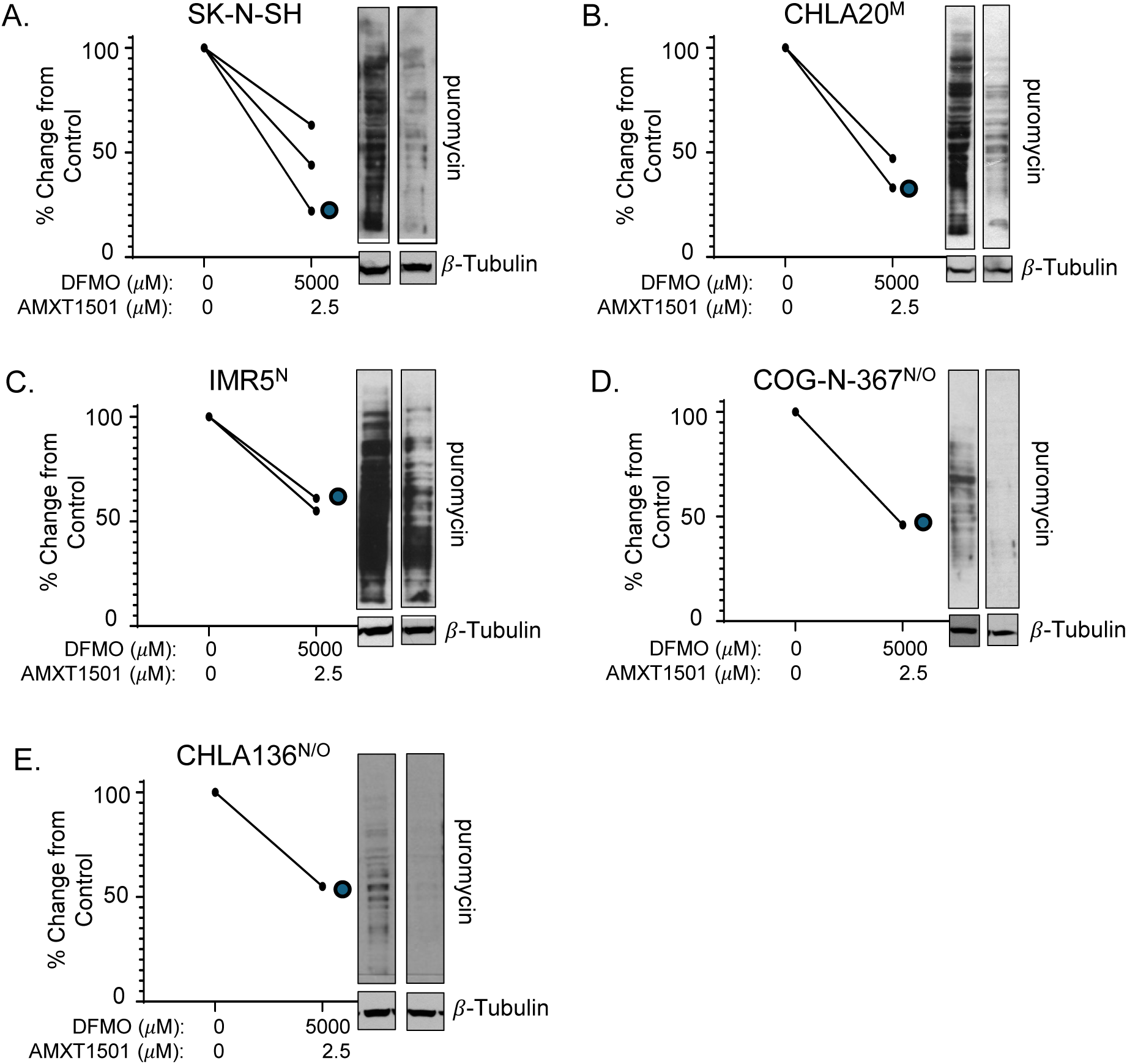
Impact of AMXT-1501 and DFMO on puromycin incorporation. Puromycin incorporation was assessed across neuroblastoma cell lines of distinct *MYCN* and *ODC1* gene status, as denoted, treated with both DFMO and AMXT-1501. Densitometry was used to define relative change from vehicle control lanes; replicates are shown in line graphs with the immunoblots corresponding to the data from the cirlce-marked line. Cell line genomic status: ^M^, *MYC* amplification; ^N^, *MYCN* amplification; ^O^, *ODC1* amplification.

### Concentration-dependent effects of DFMO on colony formation

We next assessed the impact of polyamine depletion on colony formation. As with cell viability assays [27], supratherapeutic DFMO (5 mM) significantly reduced colony formation (**Fig.6**). At therapeutically relevant DFMO concentrations up to 500 µM, effects on colony formation were modest. Notably, CHLA136 cells with co-amplification of *MYCN* and *ODC1* were highly sensitive to DFMO with marked inhibition at 150 µM, despite being relatively resistant to hypusination inhibition by DFMO.

**Figure 6.**
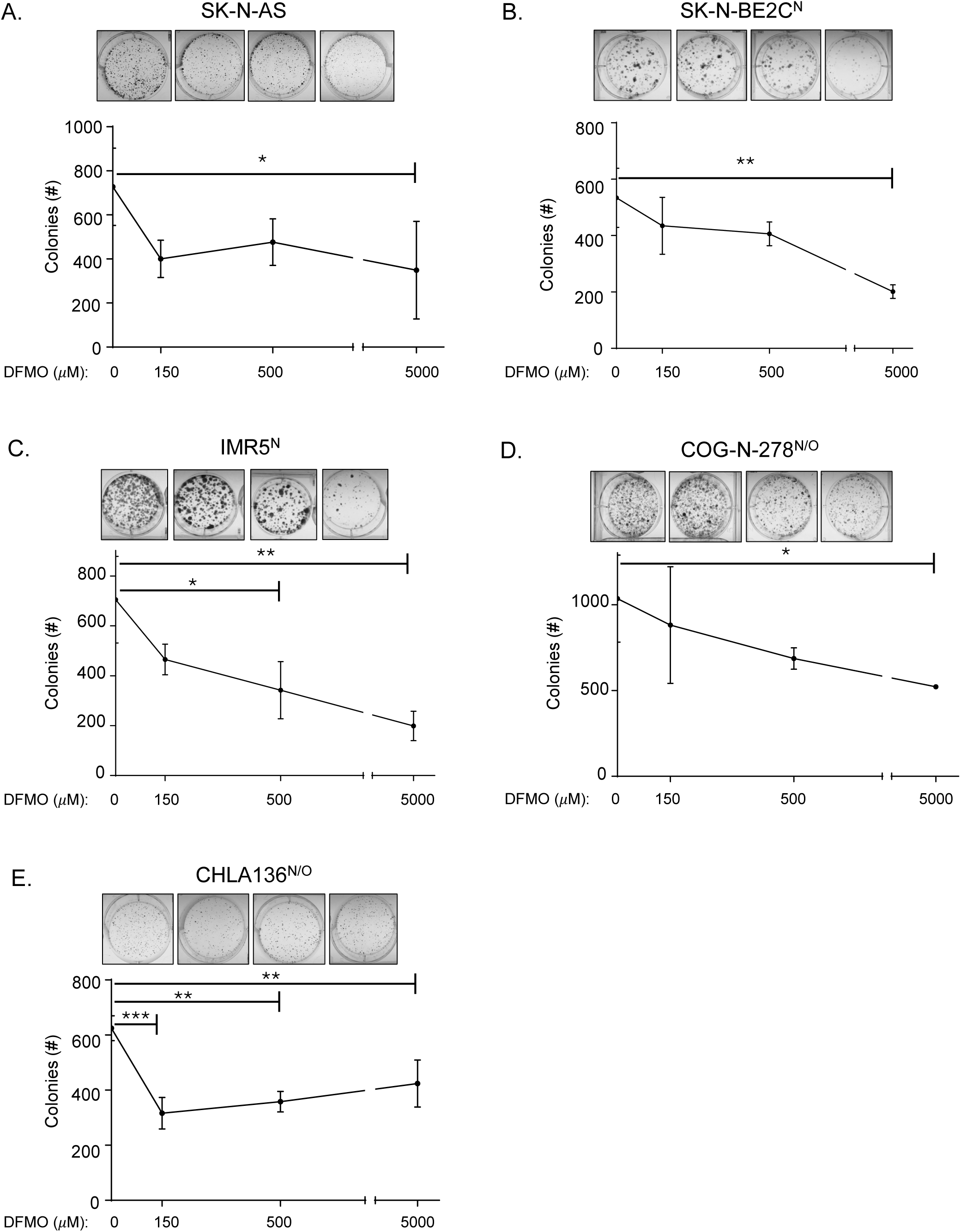
Colony Formation Assay across a range of DFMO exposures. Colony formation was assessed across neuroblastoma cell lines of distinct *MYCN* and *ODC1* gene status, as denoted, treated with DFMO across a range of concentrations. Data represents replicate wells with representative well-images shown above. Cell line genomic status: ^M^, *MYC* amplification; ^N^, *MYCN* amplification; ^O^, *ODC1* amplification; *= p<0.05, **=p<0.01, ***=p<0.001.

### DFMO treatment leads to increased ODC protein

We assessed ODC by immunoblot in cell lines with and without DFMO treatment, assessing for differential protein abundance correlated with genomic changes. We did not see marked differences in basal ODC expression across cell lines with *MYCN* amplification, *MYCN* and *ODC1* amplification, or no amplification, potentially due ODC being strongly transcriptionally regulated by MYC, which is pervasively overexpressed in neuroblastoma cell lines. We noted consistently increased ODC protein upon DFMO exposure (**Fig.7**), least pronounced in CHLA136 cells with *MYCN/ODC1* co-amplification, with relatively high ODC basal expression. However, cells with less basal ODC had clear upregulation even at lower DFMO concentrations. We hypothesize this increase may be due to DFMO being covalently linked to ODC, stabilizing the protein and impeding proteasomal degradation (ODC is degraded independent of ubiquitination; [72]). Since ODC increase occurs at clinically relevant concentrations of DFMO, it provides a potential biomarker for DFMO target engagement.

**Figure 7.**
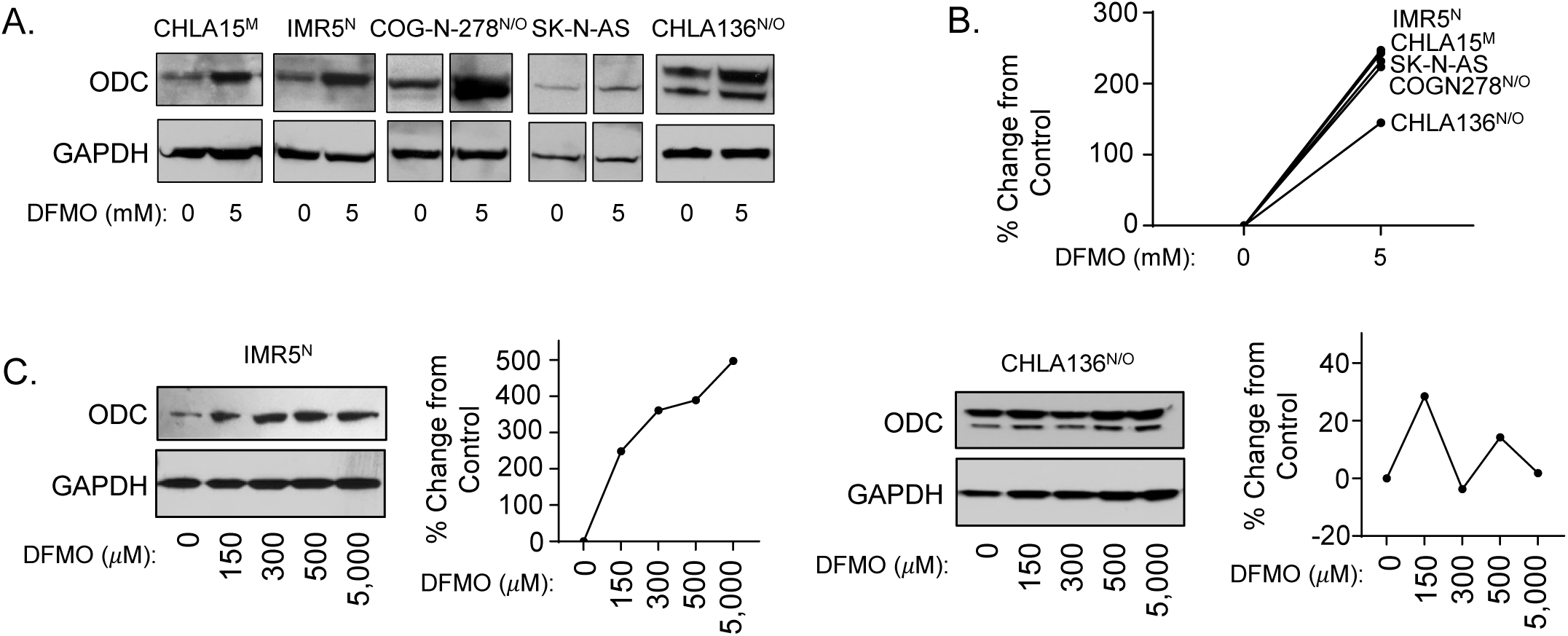
ODC protein in response to DFMO exposure. (A, B) Treatment with DFMO leads to increased ODC protein levels by immunoblot across cell lines. (C) Dose response of IMR5 and CHLA136 cells to DFMO: ODC is increased at baseline in the *ODC1-*amplified cell line without significant increase in response to DFMO; shown as immunoblot and by densitometry relative to control lanes. Cell line genomic status: ^M^, *MYC* amplification; ^N^, *MYCN* amplification; ^O^, *ODC1* amplification.

## DISCUSSION

Neuroblastoma is a lethal childhood cancer in which deregulated expression of *MYC* genes drive neoplasia-enabling transcriptional programs [4]. *MYCN* amplification is an initiating event (truncal mutation), underscoring its value as a therapeutic target [51, 73], yet MYC proteins remain difficult to target pharmacologically. An alternative to direct inhibition is to target essential oncogenic pathways downstream of MYC. Polyamine sufficiency supports protein translation and cellular biomass creation in rapidly-dividing cancers [11], is regulated by MYC [22], and represents a potential therapeutic vulnerability [74]. Indeed, DFMO at higher exposures has demonstrated anti-tumor activity in complementary murine models [22, 27, 38, 59] and is tolerable with evidence for clinical benefit in combination with chemotherapy for children with relapsed/refractory neuroblastoma [32]. However, lower dose DFMO has also been studied, largely in the chemoprevention setting for individuals at-risk for epithelial cancers [75, 76], but also in neuroblastoma patients as a post-therapy maintenance [35–37]. Here, we assessed a range of DFMO exposures recapitulating low to high-dose DFMO to assess relative anti-tumor activity and explore potential biomarkers.

In a fully penetrant and rapidly lethal transgenic neuroblastoma-prone mouse model (*TH-MYCN*^+/+^ model), DFMO monotherapy inhibited tumor progression and prolonged survival when administered at a human equivalent dose of ∼6.6 gm/m^2^/day, but not at exposures equivalent to 3.3 gm/m^2^/day or less, while the approved DFMO dose for use in neuroblastoma patients during maintenance is ≤1.5 gm/m^2^/day. We sought biomarkers of anti-tumor activity that could be elicited by high-dose DFMO and focused on protein translation as gene expression studies across diverse tumors, including neuroblastoma [8, 10], identified a role for MYC and polyamines in translation by regulating ribosomal proteins, rRNAs, tRNAs, and initiation and elongation factors (reviewed in [12]). Studies using cell-free translation systems show that polyamines stimulate translation up to 8-fold [64], although whether this effects all mRNAs or a subset (a polyamine-dependent translatome [77]) is unresolved.

Regulation of eIF4E activity through the inhibition of 4EBP1 phosphorylation has been implicated in polyamine-supported translation downstream of mTOR [67]. We were unable to elicit loss of 4EBP1 phosphorylation through supratherapeutic DFMO exposures *in vitro* suggesting this is not a mechanism of translation inhibition by polyamine depletion. We next looked at eIF5A, the sole mammalian protein hypusinated by the conjugation of spermidine with a lysine residue via deoxyhypusine synthase and deoxyhypusine hydroxylase [19]. eIF5A was initially characterized as a translation initiation factor but its function has been extended to translation elongation where it is plays a crucial role allowing the ribosome to decode polyproline and related polypeptide motifs [68–70]. Hypusination is essential for this activity and is entirely dependent on available spermidine. DFMO inhibited eIF5A hypusination at concentrations of 300-500 µM, potentially achievable with high-dose DFMO in children, in a subset of cell lines. Yet even at these exposures a large fraction of eIF5A remained hypusinated, and for many cell lines no change was elicited except at supratherapeutic DFMO exposures.

Since polyamines have also been linked to protein translation independent of 4EBP1 and eIF5A (via binding with rRNAs, tRNAs and other translation factors) we also measured puromycin incorporation into nascent polypeptide chains as a surrogate for translation. Again, effects were variable among cell lines and concentration dependent. Two of the three more sensitive cell lines (with puromycin incorporation reduced to <50% of control cells by DFMO in at least one biological replicate [IMR5, SK-N-BE2C] were also most sensitive to DFMO-mediated inhibition of eIF5A hypusination. Those least sensitive to eIF5A hypusination were among the least sensitive to puromycin incorporation inhibition. Only in SK-N-AS cells was protein translation reduced >50% at DFMO exposures of 150 µM that are readily achievable in the clinic with high-dose DFMO. Tumor and normal cells can increase their uptake of polyamines when polyamine synthesis is inhibited, and selective polyamine transporters like AMXT-1501 have been developed to block this rescue mechanism. Co-exposure of cells to DFMO and AMXT-1501 led to enhanced inhibition of protein synthesis even in cell lines that were resistant to high-dose DFMO monotherapy (CHLA20, COG-N-278). Of note, the combination of DFMO and AMXT-1501 is currently being studied in clinical trials. Finally, we found the impact of DFMO on colony formation to be concentration dependent and most pronounced at exposures above those achievable in the clinic (5 mM). The impact of 150 µM DFMO, achievable with high-dose but not low-dose DFMO, was more modest.

In this work, neither survival in mouse models nor any assessed biomarker of protein translation was altered by DFMO exposures achieved with low-dose DFMO. However, while high-dose DFMO significantly extended neuroblastoma survival in the *TH-MYCN* model, the concentrations routinely achieved also had modest effects on these same translation parameters *in vitro*. Specifically, eIF5A hypusination could be partially inhibited by DFMO and potentially impact elongation and ribosome decoding of polyproline motifs. As ∼25% of human protein isoforms harbor at least one polyproline motif (3 or more consecutive prolines [78]), disabling eIF5A might reduce measures of global translation. Indeed, the neuroblastoma cell lines with the greatest inhibition of translation as measured by puromycin incorporation were those most sensitive to eIF5A effects. We also sought correlations among biomarkers of DFMO activity among neuroblastoma cell lines differing in *MYCN* and *ODC1* copy number. We identified *ODC1* co-amplification in ∼13% of *MYCN* amplified primary tumors and cell lines and included non-amplified, *MYCN* amplified and *MYCN/ODC1* dual-amplified cell lines in this work. We did not identify any biomarker that correlated strongly with genomic status, suggesting a common vulnerability to DFMO effects or that the major effects elicited by DFMO were not studied herein. These findings are largely consistent with those of Gandra et al [79]. One additional finding of interest was that ODC protein appeared to be stabilized by DFMO *in vitro*, suggesting that this might be assessed a biomarker of target-hitting (covalent ODC protein binding by DFMO).

Limitations of this work include that tissue culture may not recapitulate *in vivo* polyamine metabolism sufficiently, as polyamine concentrations change in response to DFMO concentration, time of exposure, and availability of ODC substrate and polyamines in the media. Cell lines have also adapted to media without supplemented ornithine, the precursor of polyamines. We did not measure polyamines so cannot directly compare the intracellular levels induced with different DFMO concentrations. It is important to note that *in vitro* DFMO concentration is consistent over time, while *in vivo* BID (low-dose schedule) or TID (typical high-dose schedule) oral dosing leads to variations in concentration ranging 2- to 3-fold between C_min_ and C_max_, which is not modeled herein. Finally, we assessed candidate translation biomarkers posited to be downstream of polyamine depletion, but it is likely that a more detailed understanding of polyamine depletion effects on translation will require unbiased approaches to interrogate codon-level translation and ribosome activities, as supported recently [80].

---

**Supplemental Figure 1.**
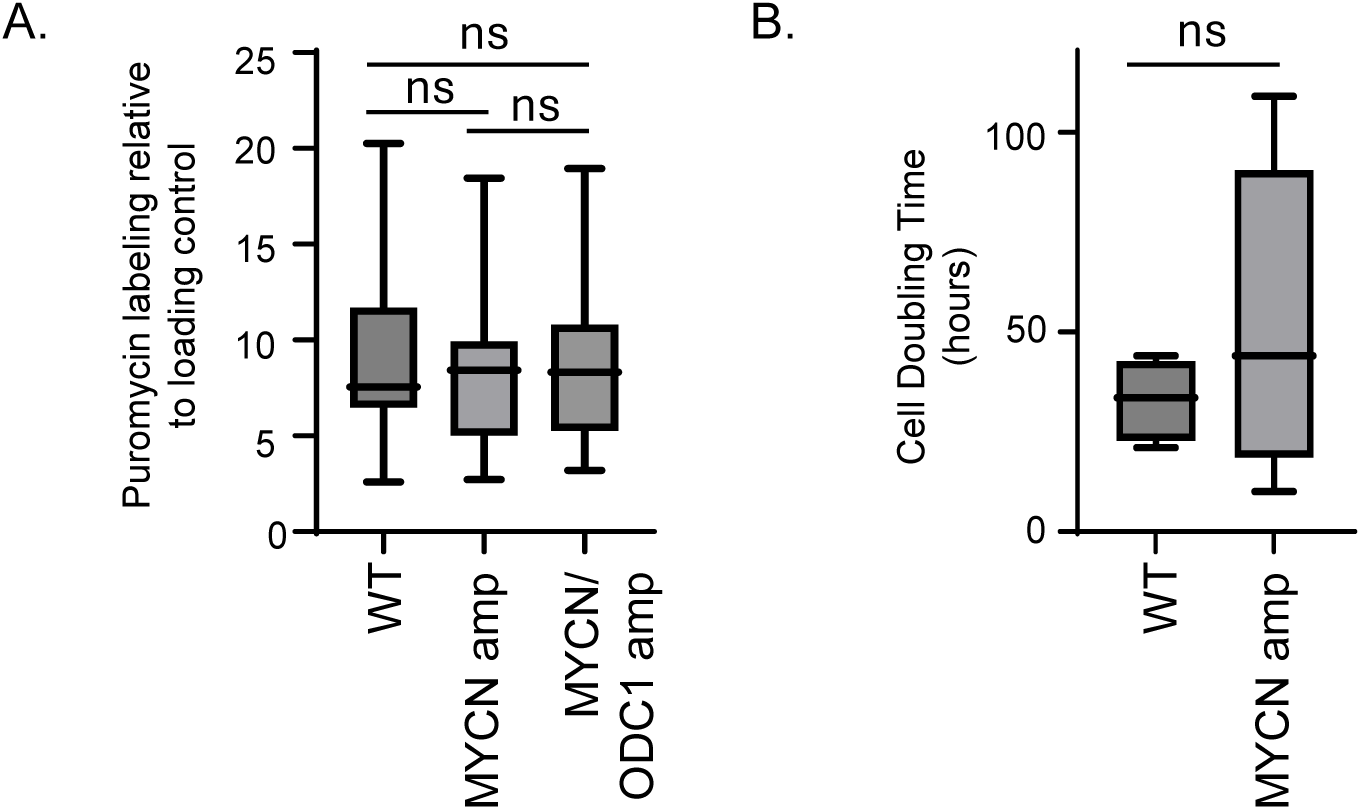
Cell line summary data by genomic status. (A) Puromycin incorporation compared to loading control via densitometry comparing non-amplified, *MYCN-*amplified, and *MYCN/ODC1* co-amplified neuroblastoma cell lines (ANOVA with multiple comparisons). (B) *In vitro* cell doubling time of non-amplified and *MYCN-*amplified cell lines (Welsh’s t-test).; ns= p>0.05.

**Supplemental Table 1.** qPCR copy number results for neuroblastoma cell lines.

| Cell Line | MYCN amplification status by qPCR | ODC1 amplification status by qPCR | Confirmation available from publication or public database [Refs. 52-54] |
| --- | --- | --- | --- |
| CHLA122 | Yes | Yes | No |
| CHLA136 | Yes | Yes | Yes |
| CHLA20 | No | No | No |
| CHLA42 | No | No | No |
| CHLA79 | No | No | No |
| CHP134 | Yes | No | Yes |
| CHP901 | Yes | No | No |
| COG-N-278 | Yes | Yes | Yes |
| IMR5 | Yes | No | Yes |
| LA-N-5 | Yes | No | Yes |
| LA-N-6 | No | No | Yes |
| NB1643 | Yes | No | Yes |
| NB69 | No | No | Yes |
| NBLS | No | No | Yes |
| NGP | Yes | No | Yes |
| NLF | Yes | No | Yes |
| SK-N-AS | No | No | Yes |
| SK-N-BE(2)C | Yes | No | Yes |
| SK-N-FI | No | No | Yes |
| SK-N-SH | No | No | Yes |
| SMS-KCN | Yes | Yes | Yes |
| SMS-KCNR | Yes | Yes | Yes |
| SMS-SAN | Yes | No | Yes |

**Supplemental Table 2.**
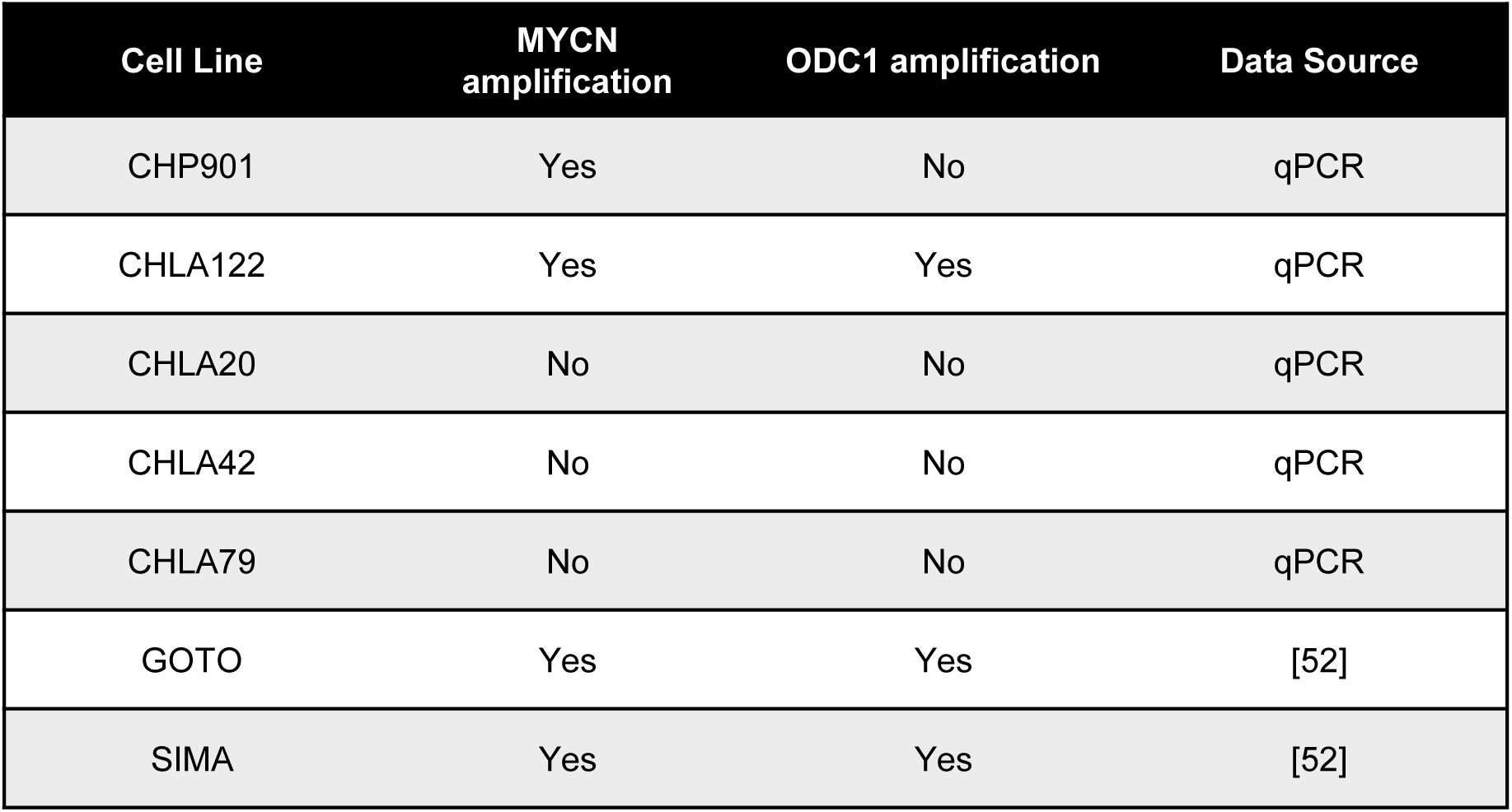
*MYCN* and *ODC*1 copy number results for neuroblastoma cell lines.

## Notes

### Competing Interest Statement

The authors have declared no competing interest.

